# Host Cellular Remodeling, Not Viral Transcriptional Inhibition, Drives Quorum Sensing Signal-Mediated Evasion of Viral Lysis in *Gephyrocapsa huxleyi*

**DOI:** 10.64898/2026.09.30.755861

**Authors:** Amanda J. Platt, Bárbara D. Bitarello, Vitor R. C. Aguiar, Natalie Goeler-Slough, Marta Wilbrink, Jiahe Wu, Emma Sijia Yu, Kristen E. Whalen

## Abstract

Bacterial infochemicals can profoundly alter phytoplankton physiology, yet how these compounds influence viral infection outcomes remains poorly understood. The bacterial quorum sensing molecule 2-heptyl-4-quinolone (HHQ) enables the globally abundant coccolithophore, *Gephyrocapsa huxleyi*, to evade virus-induced mortality following infection by *Emiliania huxleyi* virus (EhV). HHQ reduced infectious virus production at 24 hours despite largely intact viral gene transcription, as revealed by RNA sequencing of host and viral responses during EhV-201 infection. Host transcriptomics instead revealed rapid and progressive HHQ-dependent physiological remodeling evidenced by changes in protein encoding genes associated with vesicular trafficking, and cytoskeletal organization. Transcriptional responses further implicated endolysosomal sorting, membrane remodeling, and protein modification as potential barriers to productive infection. These findings indicate that HHQ-mediated evasion of virus-induced mortality occurs downstream of viral entry and transcription, through remodeling of host processes required for virion assembly, maturation, trafficking, and release.

**IMPORTANCE:** Viruses are major drivers of phytoplankton mortality in the ocean, influencing the fate of photosynthetic biomass and the movement of carbon and nutrients through marine ecosystems. Yet, the extent to which bacterial infochemicals can mechanistically alter the outcome of phytoplankton viral infections remains poorly understood. Here, we show that the bacterial signaling molecule 2-heptyl-4-quinolone alters infection of the globally important marine alga, *Gephyrocapsa huxleyi*. Rather than broadly suppressing viral gene transcription, the bacterial signal rapidly alters host processes involved in intracellular transport, protein quality control, and cytoskeletal organization. These changes may prevent infected cells from efficiently assembling, transporting, or releasing infectious viral progeny. These findings identify host physiological remodeling as a mechanism by which a bacterial chemical signal can alter the outcome of algal– virus interactions, highlighting an underappreciated role for bacterial metabolites in shaping marine viral infection and potentially the fate of phytoplankton populations.

## INTRODUCTION

Marine phytoplankton are responsible for nearly 50% of global primary production, driving the biological carbon pump, and influencing global climate by drawing down atmospheric carbon dioxide (1). Marine viruses are key modulators of phytoplankton community dynamics and productivity (2, 3) as viral infections and subsequent host cell lysis result in the turnover of approximately 20% of the oceanic photosynthetic biomass daily (4, 5). Viral-mediated phytoplankton lysis releases dissolved organic material that fuels remineralization in the microbial loop, redirects carbon from higher trophic levels (6), and influences marine community composition, succession, and global biogeochemical cycles (4, 7, 8). While abiotic factors, such as temperature, salinity, nutrients, and light/UV, are known to affect virus-phytoplankton interactions (9, 10), the influence of biotic interactions on infection outcomes and virus-host population dynamics remains poorly understood (11). Determining the full scope of how bacterial community members influence viral infection dynamics is critical for predicting phytoplankton bloom trajectories and understanding the flow of carbon and nutrients through marine ecosystems.

*Gephyrocapsa huxleyi* (formerly known as *Emiliania huxleyi*) is a globally distributed single-celled photosynthetic alga that forms the base of marine food webs and plays a key role in global carbon cycling through biomineralization of their calcite-based coccoliths (12, 13).

Massive seasonal blooms of *G. huxleyi* covering thousands of square kilometers (14) are frequently terminated by species-specific *Nucleocytoviricota* viruses (NCVs), termed *E. huxleyi* viruses (EhV), formerly classified as nucleocytoplasmic large DNA viruses (NCLDVs) (15–18). Algal EhVs belong to the family Phycodnaviridae where more than 20 viral representatives are known to infect *G. huxleyi* (15, 19–21).

Bacterial infochemicals also exert potent control over eukaryotic phytoplankton physiology (22–25), including inhibition of virus production (26) and interference with virus-mediated host processes (27). We previously described how a bacterial quorum sensing signal, 2-heptyl-4-quinolone (HHQ), disrupts DNA replication and repair machinery in *G. huxleyi*, inhibiting cell cycle progression and preventing virus-induced morality (26, 27). Because viral replication depends on commandeering the host’s replication machinery early in EhV infection (28) and HHQ disrupts key enzymes involved in nucleotide biosynthesis, HHQ treatment was hypothesized to prevent viral transcription. Our results show that viral transcription in HHQ-treated cells closely mirrors controls and is not inhibited by HHQ. Yet, HHQ treatment significantly reduced infectious virus concentrations compared with paired controls, indicating that HHQ-exposed cells produce fewer infectious virions. Analysis of host differential gene expression revealed rapid and progressive HHQ-dependent changes in cellular processes that support multiple stages of productive EhV infection. Together, these results suggest that HHQ-mediated evasion of viral-induced lysis results from host physiological remodeling and establishes how a bacterial quorum sensing signal can rapidly alter algal physiology and, in turn, reshape host-virus dynamics.

## MATERIALS AND METHODS

### Cultivation and enumeration of *G. huxleyi* and EhV-201

All experiments used axenic, non-lith-forming *G. huxleyi* CCMP2090 (National Center for Marine Algae and Microbiota). Cultures were grown in 0.2 *_μ_*m filtered, autoclaved seawater supplemented with f/2 nutrients without silica (f/2-Si) (29) under a 14-hr light (80 ± 5 *_μ_*mol photons m^-2^ s^-1^)/10 hr dark cycle at 18℃ and 35 salinity. Cultures were transferred weekly to maintain exponential growth. *E. huxleyi* virus (EhV-201), provided by Kay Bidle (Rutgers University), was propagated weekly until CCMP2090 cultures lysed to <1000 cells mL^-1^.Viral lysates were stored at 4℃ in the dark until use. *G. huxleyi* cells were enumerated live by flow cytometry (Guava, EasyCyte BGHT) using species-specific settings forward scatter, side scatter, and red fluorescence (695/50 nm) settings at 0.59 *_μ_*L s^-1^ for 3 min. Viral particles were enumerated via flow cytometry following fixation and SYBR green staining (see Supplemental Methods). Infectious virions were quantified by most probable number (MPN) assay (30) in concentrated viral lysates and samples collected after dosing each infected bottle across five trials (see Supplemental Methods).

### RNA isolation and sequencing

To examine HHQ effects on host and viral transcription, five sequential infection trials were conducted with paired non-infected controls. For each trial, an exponential-phase axenic *G. huxleyi* culture was used to inoculate four 1.8-L cultures at approximately 1.9x10^5^ cells mL^-1^. After 48 hr of growth to approximately 5x10^5^ cells mL^-1^, cultures received 100 ng mL^-1^ of HHQ or a vehicle (0.05 % DMSO) control and EhV-201 at a 1:1 host cell-to-virion ratio or a matched volume of f/2-Si, yielding four treatments (HHQ control, HHQ infected, DMSO control, DMSO infected). Subsamples (400 mL) were collected at 45 min (T1), 3 hr (T2), 8 hr (T3), and 24 hr (T4) post treatment/infection. *G. huxleyi* cells were pelleted and flash frozen in liquid nitrogen until RNA extraction (see Supplemental Methods).

### Transcriptomic analyses

Scripts to reproduce all analyses in this section are available in https://github.com/bitarellolab/Ghux_Ehv201. Sequence reads were trimmed with fastp (v0.24) (31) and unpaired reads were discarded. Paired-end reads that passed QC were mapped to a decoy-aware combined transcriptome (32) of *G. huxleyi* (GCA_000372725.1) and EhV-201 (GCA_010974645.1) and quantified using Salmon (v1.10.3) (33). DESeq2 (1.49.2) (34) was used for differential gene expression analyses excluding four samples: one sample after initial QC (HHQ_cntl_t1_3), two after exploratory data analyses (DMSO_cntl_t2_1 and, DMSO_inf_t2_1), and a final sample (DMSO_cntl_t1_1) resulting from a cracked collection bottle during centrifugation. P-values were adjusted for multiple-testing using the Independent Hypothesis Weighting (IHW) procedure (35). Host differentially expressed genes (DEGs) were defined as those with *P_IHW_*<0.05 and an absolute shrunken (36) log2-fold change in expression (*log2FC*, for simplicity) of |*log2FC*|>2, unless noted otherwise for GO enrichment analyses or comparison to previously published studies. Because few viral DEGs met these criteria, likely due in part to limited statistical power, a more permissive significance threshold (simply *P_IHW_*<0.05) was used for downstream analyses (see Supplemental Methods).

Uniprot and gene ontology (GO) annotations for *G. huxleyi* and EhV-201 were downloaded from Uniprot (release 18-Jun-2025), including KEGG annotations for *G. huxleyi.* GO enrichment was performed separately for upregulated (*log2FC>1, P_IHW_*<0.05) and downregulated (*log2FC*<-1, *P_IHW_*<0.05) DEGs from HHQ control vs DMSO control and HHQ infected vs DMSO infected comparisons at each time point using g:Profiler (37). *P* values were adjusted using the g:SCS method (38), with a significance cutoff of *P_SCS_*<0.05.

## RESULTS & DISCUSSION

### HHQ treatment reduces the amount of infectious virus in infected *G. huxleyi* cultures

To determine how HHQ affects EhV infection dynamics, HHQ- and DMSO-treated *G. huxleyi* cultures were infected with EhV-201 in a 1:1 virus-to-host ratio and monitored for host abundance and infectious viral titer for 24 hours (**Figure 1**). Consistent with previous findings (27), HHQ arrested *G. huxleyi* growth regardless of infection status (**Figure 1b**). While the amount of infectious virus increased in DMSO infected cultures from 45 min to 24 hr, the increase in HHQ treated infected cultures was not significant (two-way ANOVA, Tukey’s post-hoc test, adjusted *P* =0.9920; **Figure 1c**). The amount of infectious virus was also significantly lower in HHQ-treated cultures compared to DMSO-treated cultures at 24 hours (two-way ANOVA, Tukey’s post-hoc test, adjusted *P* = 0.0185; **Figure 1c**). Consistent with reduced infectious virion concentrations and burst sizes reported previously (26), these results suggest that HHQ impairs the formation of fully infectious progeny virions, either by reducing virion release or by disrupting processes required for proper virion assembly and maturation.

**Figure 1.**
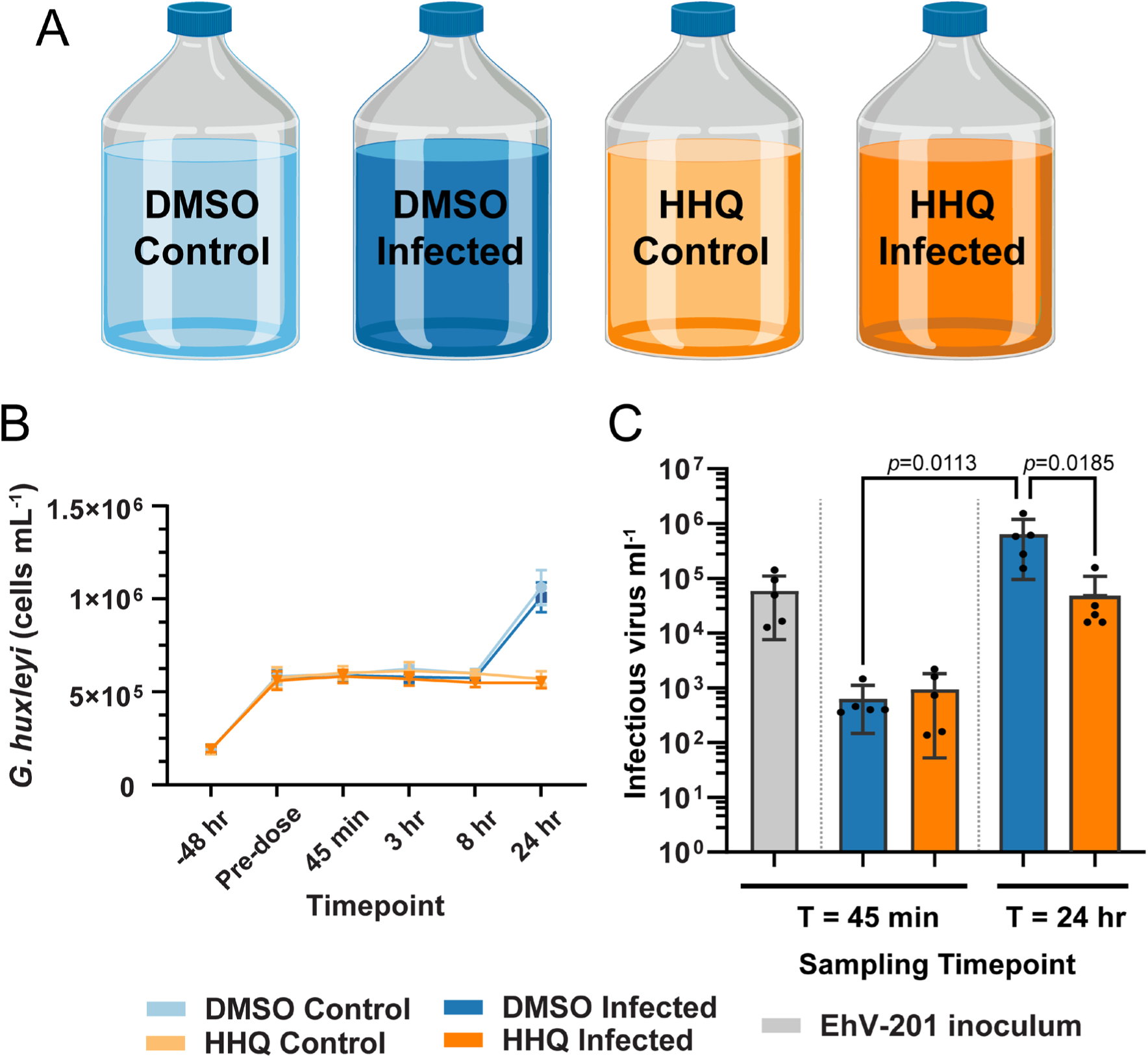
HHQ arrests *G. huxleyi* cell division and reduces infectious EhV-201 production within 24 hr. (**A**) Experimental design. Exponentially growing *G. huxleyi* cultures (1.8 L) were treated with DMSO (vehicle control) or 100 ng mL^-1^ HHQ and immediately infected with EhV-201 at a 1:1 host-to-virus ratio. Uninfected controls were maintained in parallel. Samples were collected at 45 min, 3 hr, 8 hr, and 24 hr post-infection across five biological replicates. (**B**) *G. huxleyi* cell abundance before treatment (-48 hr) and during 24 hr of infection. (**C**) Infectious virions quantified by Most Probable Number analysis at 45 min and 24 hr post-infection. Data in B and C represent mean ± SD, *n* = 5. Significance was determined by two-way ANOVA with Tukey’s post-hoc test. Adjusted *P* values are shown.

### RNA-seq reveals distinct temporal and HHQ-dependent transcriptional programs

To characterize *G. huxleyi* transcriptional responses to HHQ with and without EhV-201 infection, host and viral gene expression was profiled at 45 min (T1), 3 hr (T2), 8 hr (T3), and 24 hr (T4), capturing early molecular responses potentially contributing to HHQ-mediated evasion of virus-induced mortality. Reads were jointly aligned to the *G. huxleyi* (GCA_000372725.1) and EhV-201 (GCA_010974645.1) transcriptomes. After removal of four samples during the quality control stage (see Supplemental Methods), 76 transcriptomes yielded 16.82-41.87 million paired-end reads per sample (median 26.81 million). The majority of reads mapped to *G. huxleyi* (88.4% - 100%, median 100%), including infected samples (range 88.4% - 100%, median 99.98%). EhV-201 reads were essentially absent in T1 (45 min), but increased from median abundances of 0.015% and 0.025% at T2 and T3, respectively, to 0.84-11.6% (median 3.27%) at T4. For this reason, T1 was excluded from viral DEG analysis. Of 37,324 annotated host genes, 28,082 passed expression filtering (≥10 reads in at least four samples and ≥ 15 reads across all samples) and were retained for analysis. Of the 447 viral genes, 228 passed expression filtering after removal of T1. Gene expression distributions were highly consistent across all treatments, infection states, and time points (**Supplemental Figure 1**), indicating minimal technical variation. Differential expression was assessed across 16 pairwise comparisons to characterize HHQ and infection effects on the host (**Supplemental File 1**).

Principal component analysis identified sampling time and HHQ exposure as major sources of host transcriptomic variation (**Supplemental Figure 2**). PC1 (40% of the variance) was driven by time, with 8 hr samples clustering separately regardless of treatment or infection status (**Supplemental Figure 2a**). This transient state, absent by 24 hr, likely reflects the endogenous dusk transition from energy acquisition toward respiration, fatty acid degradation, mitochondrial metabolism, and cell-cycle progression (39). In contrast, PC2 and PC3 were driven primarily by HHQ, separating HHQ-treated samples from DMSO controls regardless of infection (**Supplemental Figure 2b**). The overlap of infected and uninfected samples within treatments indicates that HHQ exerts a stronger transcriptome-wide effect than viral infection during the first 24 hr.

### EhV-201 transcription remains largely intact in HHQ-treated infected cultures

Harvey et al. (2023) showed EhV-201 infects HHQ- and DMSO-treated *G. huxleyi* cultures at similar rates, indicating that evasion of viral induced mortality by HHQ-treated cells occurs after viral entry (26). Because HHQ inhibits *G. huxleyi* dihydroorotate dehydrogenase (DHODH), a key enzyme in *de novo* pyrimidine biosynthesis required for RNA and DNA production (40), we hypothesized that HHQ impairs viral transcription. Differential expression of viral genes between infected HHQ-treated and infected DMSO-treated cultures was therefore assessed at 3 hr, 8 hr, and 24 hr (**Supplemental File 2**). The 45 min time point was excluded because viral transcripts were largely undetectable (**Supplemental Figure 3a**), which also drove its separation along PC1 (70% of the variance; **Supplemental Figure 3b**). PC2 and PC3 explained only 4% of variance and did not differentiate HHQ treated infected cultures from DMSO treated infected cultures, consistent with limited viral gene expression (**Supplemental Figure 3c)**. Of the 447 EhV-201 protein coding genes (228 analyzed after filtering), only 22 were differentially expressed at *P_IHW_*<0.05, nine of which have UniProt protein annotations (**Table 1; Supplemental File 3**). Notably, Major Capsid Protein (MCP; EPVG_00083), an essential capsid component and marker of later EhV-201 infection (28, 41), was upregulated by HHQ at 3 hr and 8 hr (**Table 1)**. Continued expression of this essential late-stage viral gene, together with limited transcriptome-wide differences, indicates that HHQ does not broadly disrupt viral transcription and instead supports host cell remodeling as the basis of evasion from virus-induced mortality.

**Table 1.** Annotated EhV-201 genes differentially expressed in HHQ-treated infected cultures relative to DMSO-treated infected cultures. Only viral genes with protein annotations in UniProt are shown. Values indicate *log2FC* (HHQ-treated infected cultures relative to DMSO-treated infected cultures). Blank cells indicate no significant differential expression at that time point (*_PIH_*_W_ < 0.05).

| Locus tag | Protein ID | Protein Name | $\log_2FC$ | | |
| --- | --- | --- | --- | --- | --- |
|  |  |  | 3 hr | 8 hr | 24 hr |
| EPVG_00083 | AET97971.1 | Major capsid protein | 2.31 | 1.67 |  |
| EPVG_00103 | AET97991.1 | Transcription factor S-II family protein | 0.94 |  |  |
| EPVG_00216 | AET98103.1 | CRAL-TRIO domain-containing protein | 0.78 |  |  |
| EPVG_00218 | AET98105.1 | dUTP diphosphatase (EC 3.6.1.23) | 0.52 |  |  |
| EPVG_00397 | AET98284.1 | RanBD1 domain-containing protein |  |  | -0.37 |
| EPVG_00394 | AET98281.1 | HMG box domain-containing protein |  |  | -0.43 |
| EPVG_00363 | AET98250.1 | Apple domain-containing protein | -0.62 |  |  |
| EPVG_00058 | AET97946.1 | C-type lectin domain-containing protein | -0.82 |  | -0.60 |
| EPVG_00361 | AET98248.1 | deoxyuridine 5'-triphosphate nucleotidohydrolase |  |  | -0.75 |

### HHQ triggers rapid and persistent changes in host *G. huxleyi* gene expression

To identify early transcriptional responses that may explain how HHQ-treated cells evade viral mortality, we examined host gene expression at 45 min, 3 hr, 8 hr, and 24 hr following HHQ exposure. Differential expression increased over time, with 36% of transcripts significantly altered relative to DMSO controls by 24 hr (*P_IHW_*<0.05; **Table 2**). These responses partially overlapped with our previous transcriptomic analysis of *G. huxleyi* exposed to high HHQ (100 ng mL⁻¹) for 24 hr, reported in Pollara et al. (2021) (27). Of the 5,045 genes upregulated at 24 hr in HHQ-treated cultures relative to DMSO controls (|*log2FC*|>0, *P_IHW_*<0.05,), 3,411 (67.6%) were also upregulated at 24 hr in the Pollara study (27) (**Supplemental Figure 4a**). Among these shared genes, 1,275 (37.4%) were significantly upregulated within 45 min to 3 hr of HHQ exposure, and remained elevated through 24 hr, whereas 1,115 (32.7%) were detected only at 24 hr in both studies (**Supplemental Figure 4b**). Similarly, 3,295 of the 5,076 genes downregulated at 24 hr (64.9%) overlapped with the Pollara dataset (27) (**Supplemental Figure 4c**), including 1,331 (40.3%) that were downregulated within 45 min to 3 hr and remained suppressed through 24 hr (**Supplemental Figure 4d**). Together, these results indicate that a substantial fraction of HHQ-induced host transcriptional reprogramming begins within hours of HHQ exposure and persists for at least 24 hr, independent of infection.

**Table 2.**
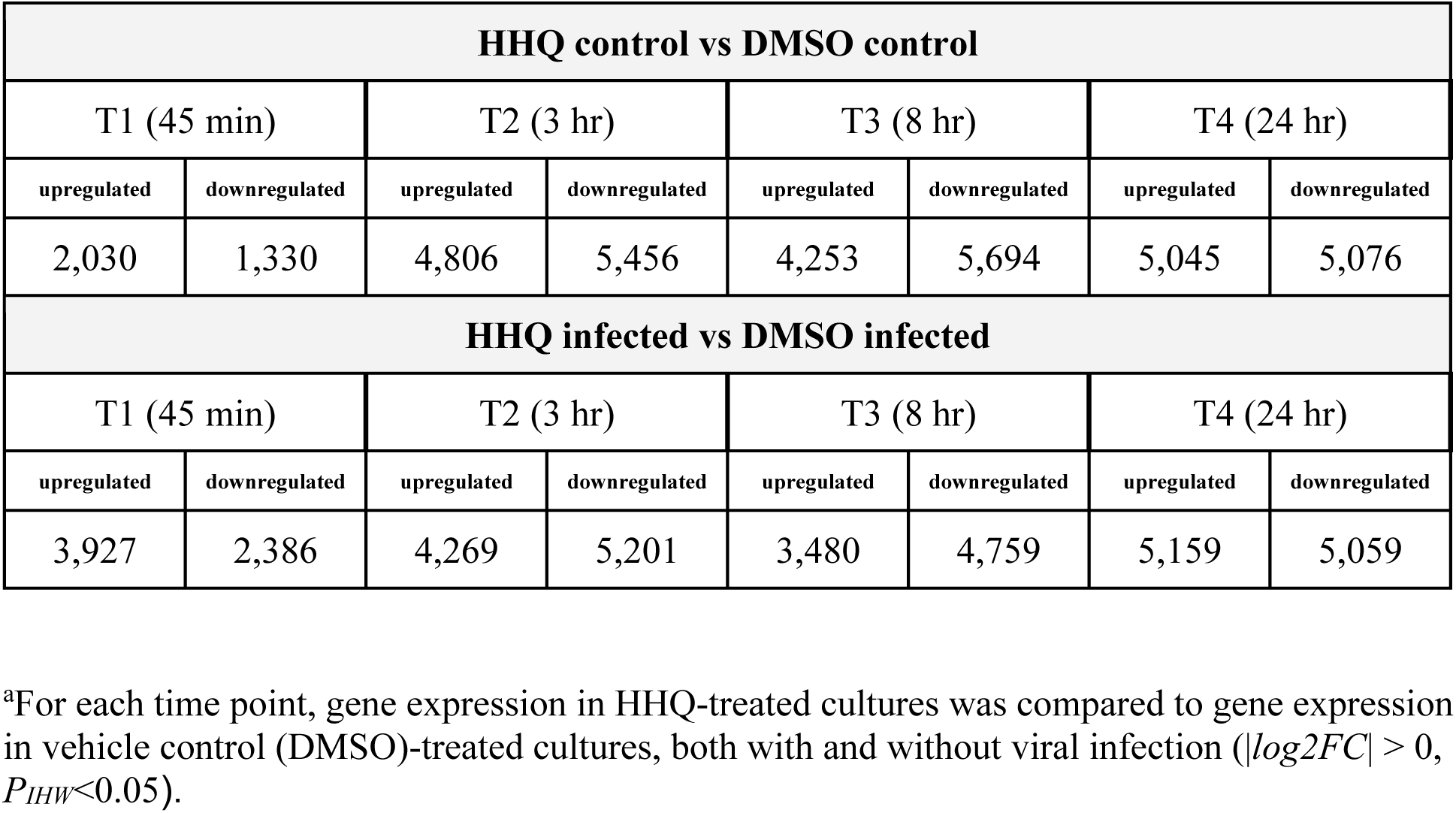
Numbers of differentially expressed genes in HHQ-treated cultures relative to DMSO controls in infected and uninfected *G. huxleyi* cultures^a^

| HHQ control vs DMSO control |  |  |  |  |  |  |  |
| --- | --- | --- | --- | --- | --- | --- | --- |
| T1 (45 min) |  | T2 (3 hr) |  | T3 (8 hr) |  | T4 (24 hr) |  |
| upregulated | downregulated | upregulated | downregulated | upregulated | downregulated | upregulated | downregulated |
| 2,030 | 1,330 | 4,806 | 5,456 | 4,253 | 5,694 | 5,045 | 5,076 |
| HHQ infected vs DMSO infected |  |  |  |  |  |  |  |
| T1 (45 min) |  | T2 (3 hr) |  | T3 (8 hr) |  | T4 (24 hr) |  |
| upregulated | downregulated | upregulated | downregulated | upregulated | downregulated | upregulated | downregulated |
| 3,927 | 2,386 | 4,269 | 5,201 | 3,480 | 4,759 | 5,159 | 5,059 |

<sup>a</sup>For each time point, gene expression in HHQ-treated cultures was compared to gene expression in vehicle control (DMSO)-treated cultures, both with and without viral infection ( $|\log_2 FC| > 0$ , $P_{IHW} < 0.05$ ).

Pollara et al. (2021) demonstrated that delaying HHQ treatment until 24 hr after EhV infection abolished evasion of virus-induced mortality (27), indicating that HHQ must rapidly reprogram host physiology to establish viral tolerance. Consequently, the earliest transcriptional responses to HHQ are likely critical for initiating this protective state. Of the 3,360 genes differentially expressed (|*log2FC*|> 0, *P_IHW_* < 0.05) at 45 min in the HHQ control vs DMSO control comparison, 1,929 were unique to this time point, whereas 358 genes remained differentially expressed through 24 hr in both the current and Pollara studies. Applying a stricter cutoff for functional classification (|*log2FC*|>1, *P_IHW_*<0.05) revealed early changes in genes encoding proteins associated with signal transduction and regulation, as well as redox metabolism and homeostasis (**Supplemental File 4**). As the first assessment of *G. huxleyi* transcriptional responses to HHQ before 24 hr, these findings suggest that rapid signaling and redox responses may initiate host physiological changes associated with evasion of virus-induced mortality.

### GO enrichment analysis reveals progressive host remodeling during HHQ-treated EhV infection

To determine which processes were shared or unique to the HHQ response in infected and uninfected cells, GO enrichment analyses were performed on differentially expressed genes (|*log2FC*|>1, *P_IHW_*<0.05,) in the Control (HHQ control vs. DMSO control) and Infected (HHQ infected vs. DMSO infected) contrasts at each time point (**Figure 2**). No GO terms were significantly enriched at 45 min, with enrichment first emerging at 3 hr and becoming more extensive through 24 hr. Many enriched GO terms were shared between the Control and Infected contrasts, indicating that HHQ treatment, rather than infection status, was the predominant driver of these host transcriptional responses. At 3 hr, upregulated genes in both contrasts were enriched for the Molecular Function term structural molecule activity, while downregulated genes were enriched for the Cellular Component term cytoplasm. Remaining categories, nucleobase-containing compound biosynthetic process and intracellular protein-containing complex, were uniquely enriched in the control comparison only. By 8 hr and 24 hr, enrichment expanded substantially and was dominated by downregulated genes. Enriched Biological Process terms shared between Control and Infected comparisons included localization, macromolecule modification, and RNA metabolic process; Molecular Function terms included catalytic activity acting on proteins or RNA; and Cellular Component terms included membrane, intracellular anatomical structure, and membrane-bounded organelle. In contrast, enrichment among upregulated genes remained limited, with structural molecule activity representing the principal shared Molecular Function term in Control and Infected comparisons.

**Figure 2.**
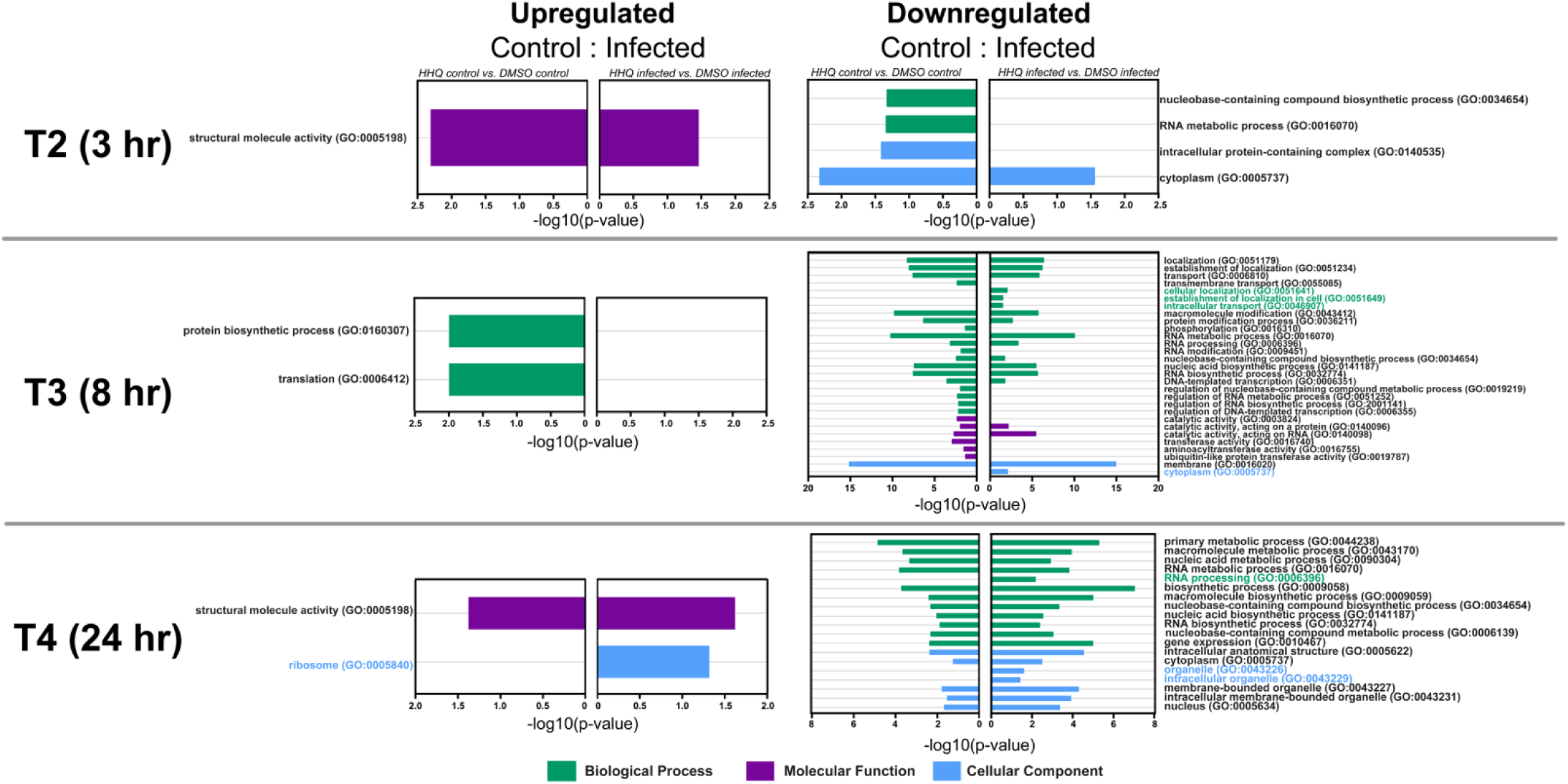
HHQ treatment is the predominant driver of host transcriptional responses. Gene Ontology enrichment analyses of HHQ control vs. DMSO control (Control) and HHQ infected vs. DMSO infected (Infected) comparisons. Upregulated (*log2FC* > 1, *P_IHW_* <0.05) and downregulated (*log2FC* < -1, *P_IHW_* <0.05) gene sets were analyzed separately at 3 hr, 8 hr, and 24 hr using g:Profiler (37). Bars show -log10(*p*-value) and are colored by GO category. Control and Infected comparisons are shown side by side. Colored GO term labels identify terms enriched exclusively in the infected comparison.

Despite substantial overlap between Control and Infected cells at 8 hr and 24 hr, four GO terms at each time point were enriched specifically in HHQ-treated infected cells but not in HHQ-treated uninfected cells (**Figure 2**). Although most intersecting genes within these categories were significantly differentially expressed (|*log2FC*|>1, *P_IHW_* <0.05) in both Infected and Control comparisons (**Supplemental File 5**), functional annotations were used to identify genes and processes in these lists that may contribute to HHQ-mediated evasion of viral mortality. At 8 hr, the broad cellular localization (GO:0051641) category included genes associated with microtubule regulation and endoplasmic reticulum (ER)-associated trafficking. The TOG domain-containing protein (EMIHUDRAFT_453318), was strongly downregulated (*log2FC*=−5.32; *P_IHW_*=2.07×10^−46^), consistent with altered microtubule dynamics given the role of TOG domains in tubulin binding and microtubule regulation (42, 43). The putative Sar1 GTPase (EMIHUDRAFT_432137) that initiates vesicle formation at the endoplasmic reticulum (44), was also downregulated (*log2FC*=−1.02; *P_IHW_*=4.16×10^−8^) (**Supplemental File 5**). These transcriptional changes may be relevant to the EhV replication cycle, which relies on nuclear and ER-derived membranes for capsid assembly and genome packaging, followed by membrane acquisition and vesicular trafficking of newly assembled virions (41).

Consistent with altered intracellular organization, the GO term cytoplasm (GO:0005737) was also enriched among downregulated genes at 8 hr. Three α-tubulin genes, (EMIHUDRAFT_104197, EMIHUDRAFT_116863, and EMIHUDRAFT_415405), were strongly downregulated (*log2FC*=−4.99, −4.06, and −4.03; *P_IHW_*=1.49×10^−22^, 3.61×10^−9^, and 1.86×10^−32^, respectively), together with the CCT/TRiC chaperonin subunits CCT-beta (EMIHUDRAFT_425512), CCT-epsilon (EMIHUDRAFT_418001), and CCT-theta (EMIHUDRAFT_109999), which contribute to tubulin folding (45). Calreticulin and calnexin, ER-resident glycoprotein chaperones (46), were similarly reduced (EMIHUDRAFT_422633; *log2FC*=−1.66; *P_IHW_*=2.81×10^−18^ and EMIHUDRAFT_49942; *log2FC*=−1.42; *P_IHW_*=2.53×10^−5^ respectively) (**Supplemental File 5**). The coordinated suppression of microtubule components, their folding machinery, and ER chaperones suggests altered cytoskeletal organization, intracellular trafficking, and ER protein-quality-control in HHQ-treated infected cells. Such changes could affect the extensive cytoplasmic and ER remodeling required for EhV assembly, maturation, and trafficking (41), potentially contributing to reduced production of infectious virions despite continued viral transcription.

Additional downregulated genes suggest effects on cellular processes that may also influence productive infection. The transcript encoding GTR1/Rag-family GTPase (EMIHUDRAFT_371787) was downregulated in HHQ-treated infected cells at 8 hr (*log2FC*=−1.04; *P_IHW_*=6.10x10^−10^). In other eukaryotes, Rag-family GTPases regulate nutrient-responsive TOR signaling at lysosomal or vacuolar membranes (47, 48), making this gene potentially relevant to the localization of EhV particles within endolysosomal compartments previously observed in HHQ-treated cells (26). Downregulation of EMIHUDRAFT_371787 may therefore reflect altered endolysosomal signaling during HHQ-treated infection, although its localization and function in *G. huxleyi* remain unknown. Finally, three light-harvesting proteins, (EMIHUDRAFT_433058, EMIHUDRAFT_435222, and EMIHUDRAFT_433847), were downregulated (*log2FC*=−1.34, −1.38, and −1.13, respectively) (**Supplemental File 5**). Vincent et al. (2021) demonstrated that EhV infection is accompanied by a progressive decline in host photosynthetic regulation, including decreased chlorophyll and loss of *psbA* expression, while photosynthetically active infected cells were associated with early virion release (49). Reduced expression of light-harvesting genes in HHQ-treated infected cells at 8 hr may therefore contribute to a cellular metabolic state less favorable for efficient viral production. Collectively, these infection-dependent transcriptional responses at 8 hr implicate cytoskeletal organization, trafficking, protein quality control in the ER, endolysosomal signaling, and photosynthetic function as host processes potentially altered by HHQ in ways that could limit productive EhV infection.

At 24 hr, enrichment of the terms organelle (GO:0043226) and intracellular organelle (GO:0043229) in HHQ-treated infected cells was driven by coordinated downregulation of genes associated with photosynthesis, cytoskeletal organization, and nuclear processes (**Figure 2**).

Photosynthetic genes were particularly prominent, including light-harvesting proteins (EMIHUDRAFT_435222; *log2FC*=−1.52; *P_IHW_*=4.55×10⁻²^6^ and EMIHUDRAFT_433058; *log2FC*=−1.69; *P_IHW_*=1.62×10⁻²²), photosystem I subunit (EMIHUDRAFT_449561; *log2FC* = −1.28; *P_IHW_*=3.07×10⁻²²), and photosystem II-associated gene (EMIHUDRAFT_428354; *log2FC* =−1.63; *P_IHW_*=8.06×10⁻²²). This broader suppression of photosynthetic machinery is consistent with Vincent et al. (2021), who observed a profound loss of host photosystem II expression during late EhV infection (49). Additionally, four histone H2A genes (EMIHUDRAFT_349615, EMIHUDRAFT_374282, EMIHUDRAFT_371985, and EMIHUDRAFT_358281) were downregulated, likely reflective of the altered chromatin organization during infection characterized by substantial nuclear remodeling and loss of normal chromatin organization (41). Finally, cytoskeletal components were also reduced, including putative actin (EMIHUDRAFT_441334; *log2FC* =−30.06; *P_IHW_*=5.39×10⁻⁷), three myosin motor-domain proteins (EMIHUDRAFT_200475, EMIHUDRAFT_76086, and EMIHUDRAFT_450146), and α-tubulin (EMIHUDRAFT_445562), suggesting altered cytoskeletal organization and intracellular transport (**Supplemental File 5**). Notably, actin (EMIHUDRAFT_441334) was not significantly differentially expressed in the Control comparison distinguishing the effects of HHQ and infection from the broader effects of HHQ treatment alone. Because microtubule- and actin-based motor systems support intracellular positioning and movement of membranes, vesicles, and organelles, their coordinated suppression could influence the extensive intracellular membrane rearrangements and trafficking associated with the EhV life cycle (41).

Together, these infection-dependent transcriptional responses reveal a progressive alteration of host cellular processes during HHQ-treated EhV infection. By 8 hr, these responses include endolysosomal signaling and photosynthetic function, while at 24 hr, broader suppression of photosynthetic machinery, cytoskeletal and motor proteins, and nuclear/chromatin-associated processes was evident. Many of these functions intersect with the extensive nuclear, ER membrane, cytoplasmic, and vesicular remodeling required for EhV genome replication, virion assembly, maturation, and release described extensively by Homola et al. (2024) (41). Thus, rather than preventing viral entry or transcription, HHQ may initiate early host physiological remodeling that progressively establishes a cellular environment less conducive to the efficient production of infectious progeny.

### Opposing transcriptional responses reveal candidate mechanisms of HHQ-mediated viral evasion

To identify robust host responses to viral infection that were altered by HHQ, we compared infection-induced differential expression in HHQ-treated cells (HHQ infected vs. HHQ control) with that of DMSO-treated cells (DMSO infected vs. DMSO control). This analysis identified protein-coding genes that responded significantly (|*log2FC*|>2, *P_IHW_*<0.05) to infection under both conditions but in opposing directions (**Table 3**). Given the limited number of differentially expressed genes meeting these criteria, we focused on individual gene responses rather than enrichment analysis. These divergent responses may provide insight into HHQ- mediated evasion of viral mortality, as HHQ does not prevent EhV-201 infection of the host but substantially reduces infectious virion production (26), despite continued viral transcription.

**Table 3.**
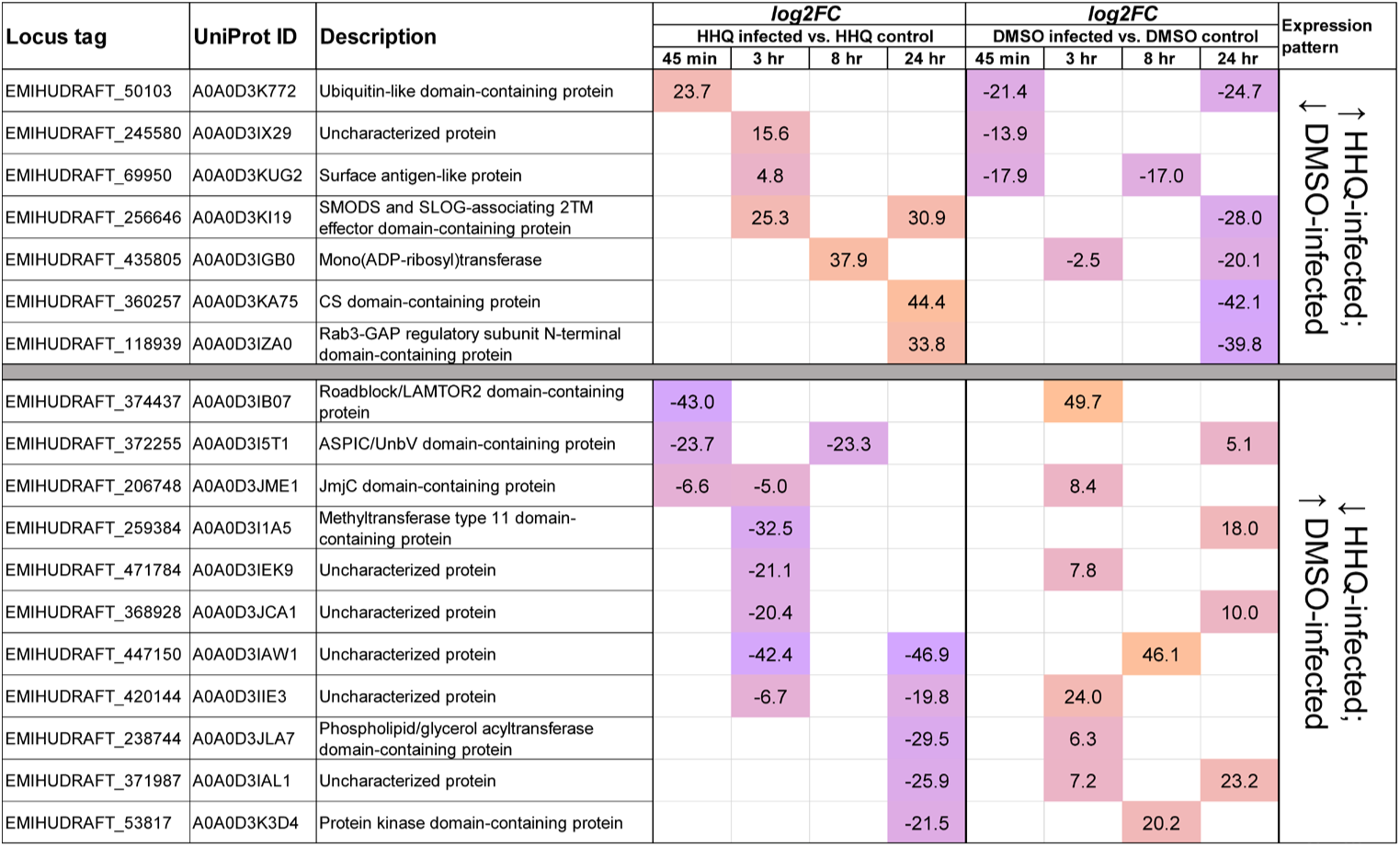
*G. huxleyi* genes with opposing expression patterns during infection under HHQ treatment versus the normal course of infection. List of genes that were differentially expressed (|l*og2FC*| >2 and *P_IHW_* <0.05) in both HHQ-infected vs. HHQ-control and DMSO- infected vs. DMSO-control comparisons, but in opposite directions. Blank cells indicate no significant differential expression at that time point. The complete list of shared differentially expressed genes is provided in **Supplemental File 6**.

EhV-201 repurposes host ER and outer nuclear membranes into membrane segments that form a major component of viral factories and provide sites for genome packaging and capsid assembly (41). Expression of EMIHUDRAFT_238744, which encodes for an ER-associated phospholipid/glycerol acyltransferase domain-containing protein, was downregulated in HHQ- infected cells at 24 hr (*log2FC*=−29.5) but induced during DMSO-treated infection at 3 hr (*log2FC*=6.3). Glycerolipid acyltransferases contribute to phosphatidic acid and glycerophospholipid synthesis and can influence cellular membrane composition (50). Previous studies have demonstrated extensive reprogramming of host lipid metabolism during productive EhV infection, including changes in fatty acid, glycerolipid, and sphingolipid metabolism that support viral replication and assembly (51–54). Together with the identification of ER-derived membranes as sites of EhV particle assembly, these observations raise the possibility that altered regulation of EMIHUDRAFT_238744 in HHQ-treated cells may contribute to processes that disrupt host phospholipid synthesis or remodeling, producing a membrane environment less conducive to viral-factory formation and/or virion assembly.

Following genome packaging, EhV-201 particles bud into intracellular vesicles, acquiring their outer membrane before trafficking toward exocytotic release (41). Several genes associated with membrane organization and vesicular transport showed striking opposing responses that could potentially influence these later stages of infection.

EMIHUDRAFT_118939, encoding a Rab3-GAP regulatory subunit N-terminal domain-containing protein, was induced at 24 hr with HHQ (*log2FC*=33.8) but strongly suppressed with DMSO (*log2FC*=−39.8). In addition to regulating Rab3, the Rab3GAP complex functions as a guanine nucleotide exchange factor for Rab18 and is required for its localization to the ER, where Rab18 contributes to membrane organization, tethering, and ER–lipid droplet contacts (55). Rab18-dependent membrane trafficking is exploited by diverse viruses for replication, intracellular transport, and virion assembly (56–61), suggesting that altered Rab3GAP expression could modify the membrane environment used for productive EhV infection. HHQ also reversed regulation of EMIHUDRAFT_256646, encoding a SMODS- and SLOG-associating two-transmembrane (SLATT) effector domain-containing protein, which was induced with HHQ (*log2FC*=30.9) but suppressed with DMSO (*log2FC*=−28). SLATT proteins are two-transmembrane effectors associated with nucleotide-centered biological conflict and defense systems, with conserved polar residues in some family members predicted to form aqueous membrane channels that may alter membrane permeability or transport. Although their function during viral infection remains unknown, SLATT proteins also occur in nucleo-cytoplasmic large DNA viruses (62). Collectively, the opposing regulation of genes encoding a Rab3-GAP regulatory subunit, and SLATT-domain suggests that HHQ may alter the cellular membrane organization important for assembly and membrane acquisition of infectious EhV particles.

Following assembly and membrane acquisition, EhV-201 particles contained within intracellular vesicles are ultimately routed toward exocytotic release, making the intracellular fate of virus-containing compartments an important determinant of productive infection (41). A potential alteration in the intracellular fate of EhV particles is further suggested by the opposing regulation of the Roadblock/LAMTOR2 domain-containing EMIHUDRAFT_374437. This gene was strongly suppressed at 45 min in HHQ-infected cells (*log2FC*=−43) but induced during DMSO-treated infection at 3 hr (*log2FC*=49.7). LAMTOR2 is associated with late endosomal/lysosomal organization, trafficking, and signaling (63, 64). This observation also complements previous observations that apparent EhV particles accumulate within endolysosomal compartments in HHQ-treated cells, whereas particles in untreated infected cells were associated with suspected early endosomes (26). HHQ-dependent alteration of endosomal maturation or sorting could favor sequestration of EhV particles within endolysosomal compartments rather than trafficking toward pathways supporting productive infection and release. More broadly, the opposing infection response extended to two additional genes encoding predicted membrane- or cell surface-associated proteins, including the surface antigen-like EMIHUDRAFT_69950, and the ASPIC/UnbV domain-containing EMIHUDRAFT_372255. Although their specific functions are unknown, their contrasting transcriptional regulation in HHQ- versus DMSO-treated infection aligns with HHQ treatment altering host membrane and cell-surface processes during infection, changes that could contribute to the distinct intracellular fate and reduced release of EhV particles.

Finally, opposing transcriptional responses extended to protein modification and turnover. EMIHUDRAFT_435805, a multi-domain protein with predicted mono(ADP-ribosyl)transferase activity, was strongly induced with HHQ at 8 hr (*log2FC*=37.9) but suppressed during DMSO-treated infection at 3 hr and 24 hr (*log2FC*=−2.5 and −20.1). Although EMIHUDRAFT_435805 is not a poly(ADP-ribose) polymerase (PARP), its predicted activity implicates a related form of ADP-ribosylation. This finding is relevant as HHQ was previously shown to increase PARP transcript and protein abundance while inhibiting PARP enzymatic activity (27), suggesting that multiple components of ADP-ribosylation may be altered in response to HHQ. Because mono-ADP-ribosylation can alter protein activity, stability, localization, and degradation, induction of EMIHUDRAFT_435805 could modify host or viral protein functions required for productive infection (65). Similarly, the ubiquitin/ubiquitin-like domain-containing EMIHUDRAFT_50103 was strongly induced with HHQ at 45 min (*log2FC*=23.7) but suppressed with DMSO at 45 min and 24 hr (*log2FC*=−21.4 and −24.7).

Ubiquitin-dependent pathways can either support viral replication through regulation of host and viral proteins or restrict infection through their degradation or altered function (66, 67). The opposing regulation of these two genes therefore suggests that HHQ may also alter the post-translational environment encountered by EhV during infection.

Collectively, the opposing transcriptional responses distinguish the cellular response to productive EhV infection from infection in the presence of HHQ, and identify several potential points of interference downstream of viral entry and transcription. Alterations in ER membrane remodeling, vesicular transport, endolysosomal sorting, and protein modification provide candidate mechanisms by which HHQ could disrupt the assembly and release of infectious progeny.

### HHQ Rewires the Conserved Viral Resistance Network Rather Than Recapitulating the Recovered Resistant Phenotype

To assess whether HHQ-mediated evasion from viral mortality shares transcriptional features with naturally virus-resistant *G. huxleyi* strains, transcriptomic profiles were compared with the core viral resistance genes recently identified by Shaler et al. (2025) (68). Their meta-analysis identified a set of 206 genes that are consistently upregulated across all published virus-resistant strains relative to their virus-sensitive counterparts (68–71). Of the 49 resistance genes in this category that had *G. huxleyi* CCMP1516 locus tags, all but one were differentially expressed (|*log2FC*| >0.5, *P_IHW_*<0.05,) in comparisons of HHQ versus DMSO in this study regardless of infection, indicating HHQ is driving the differential expression, not infection status (**Supplemental File 7**). Using UniProt annotations (72), functional predictions could be assigned to 40 genes revealing increased expression of proteins with putative domains involved in lipid trafficking (CRAL/TRIO-domain containing protein) (73), cell signaling and regulation (WW domain-containing protein) (74), amino acid transport (amino acid transporter transmembrane domain-containing protein), redox homeostasis (thioredoxin domain-containing protein) (75), and protein metabolic processes (peptidase S1 domain-containing protein) (76) (**Supplemental File 7**). The remaining genes that were upregulated across all naturally resistant strains were downregulated or showed mixed regulation in this study (**Supplemental File 7**). Together, these findings suggest that HHQ does not fully recapitulate the transcriptional program of naturally recovered virus-resistant populations, but instead establishes a distinct physiological state that may prime cells for viral tolerance without fully inducing the resistance phenotype that emerges following recovery from infection.

## CONCLUSION

Collectively, these findings reveal that HHQ-mediated evasion from EhV mortality is driven by rapid and sustained remodeling of host physiology. Building on previous work demonstrating that HHQ does not prevent viral entry, this study revealed that EhV transcription also remains largely intact during HHQ treatment, indicating that HHQ-mediated evasion of viral mortality occurs downstream of these early stages of infection. HHQ initiates changes in membrane organization and intracellular trafficking within the earliest stages of infection, followed by broader alterations in ER proteostasis, cytoskeletal organization, photosynthesis, and endolysosomal processes that may interfere with virion assembly, maturation, trafficking, and release. These findings reveal a previously unrecognized mechanism by which a bacterially produced chemical signal alters the outcome of a eukaryotic host-virus interaction by reprogramming host cellular processes required for productive infection.

## DATA AVAILABILITY

Raw sequencing reads have been added to the Gene Expression Omnibus repository (accession GSE348739), but are not yet publicly available.

## ACKNOWLEDGMENTS

Financial support to K.E.W was provided by the Charles E. Kaufman Foundation Integrated Research Education Grant (KA2021-121932), the National Science Foundation (IOS 2041748), and the Henry Dreyfus Teacher Scholar Award.

